# From Microscale to Microbial Insights: Validating High-Throughput Microvolume Extraction Methods (HiMEx) for Microbial Ecology

**DOI:** 10.1101/2025.05.25.655987

**Authors:** Marjan Ghotbi, Mitra Ghotbi, Elisa D’Agostino, Maarten Kanitz, David M. Needham

**Affiliations:** GEOMAR Helmholtz Centre for Ocean Research Kiel, 24148 Kiel, Germany; Faculty of Mathematics and Natural Sciences, Kiel University, 24118 Kiel, Germany; Middle Tennessee State University, Murfreesboro, TN, 27127, USA

## Abstract

Extracting and directly amplifying DNA from small-volume, low-biomass samples would enable rapid, ultra-high-throughput analyses, facilitate studying microbial communities where large-volume sample collection is challenging. This can aid where ‘conventional’ filtration-based methods miss capturing smaller microbes, or where microscale variability matters, such as the ocean. Here, we develop and validate physical and chemical-based DNA extractions from microvolumes with universal rRNA gene amplicons and metagenomic sequencing of all domains and viruses, on natural seawater and experimentally manipulated marine waters. Compared to 500-mL filter-based extraction, direct PCR of 3 μL of lysed seawater samples consistently captured comparable microbial community composition and diversity, with reliable amplification and little to no contamination. Metagenomic results of 10 μL of lysed microvolume samples captured high-quality assemblies, bacterial genomes, and diversity on par or better than our conventional extraction (48 vs 44 representative MAGs), and substantially more putative complete circular viral genomes (38 vs 1). Our approach enables scaling of rRNA gene sequencing and metagenomic library prep tremendously for a fraction of the cost of conventional methods and builds upon existing microvolume approaches by removing unnecessary expenses, like excess plasticware and expensive bead clean-up. The method expands opportunities for more comprehensive microbial community monitoring and controlled laboratory experiments.

## Introduction

Molecular characterizations, both taxonomic and genomic, of microbial communities is essential for understanding microbial diversity and dynamics. Traditionally, this involves collection of substantial amounts of biomass that require significant time, effort, and resources to collect, extract, and analyse. However, if sample biomass could be scaled down to be performed in microvolumes, then sample numbers could be scaled up tremendously. Applications of such methods can include high-throughput experimentation that enables testing of numerous ecological factors within a single experiment, with high replication and greater statistical power, while also reducing costs, situations where sample volume or biomass has limitations and studies where microscale diversity is important, such as the phycosphere of microalgae [1]. Although innovative microvolume extraction methods [2, 3] have achieved success in the molecular characterization of community composition, a bottleneck, specifically for high-throughput studies (e.g., 1000s of samples), remains the relatively high cost of DNA purification. Direct amplification of extracted DNA or a minimal, yet cost-effective purification step would be a preferred alternative if high-throughput analyses are required since it excludes the expensive bead-purification or centrifugation and DNA loss during purification step. Here, we develop and evaluate different microvolume extraction methods, coupled with both amplicon and metagenomic analysis. We demonstrate that this approach facilitates high-throughput lab experiments and captures diversity across all domains and viruses, to provide insight into microbial dynamics and interactions.

For High-throughput Microvolume Extraction (HiMEx) methods, we used either physical extraction through thermal shock (i.e., freeze-thaw, FT) and/or a combination of physical and chemical extraction which included addition of the lytic enzymes alone to the physical disruption (i.e., FT + proteinaseK, FTP), or a combination of lytic enzyme and a surfactant (i.e., FTP + IGEPAL, FTPIG) [3] (Fig. S1, Supplementary Information). For testing the efficacy of the HiMEx methods, we first compared them to amplicon analysis from natural surface seawater samples (Baltic Sea) collected bimonthly at different timepoints during August to November 2022 (54°19.813’ N, 10°8.993’ E), with prokaryotic and virus-like-particle abundance ranging from 0.97 – 7.56 × 10^6^ and 0.56 – 5.04 ×10^7^ per mL, respectively (Tab. S1). Quadruplicate samples of 500 mL filtered seawater (0.2 μm) and 100, 200, 400, 1000 μL of whole seawater were collected for ‘conventional’ extraction vs HiMEx (exceptions indicated in metadata, Tab. S2), respectively. All samples were immediately stored at -80°C. For filter-based ‘conventional’ extraction, samples were extracted via DNeasy Plant Kit (QIAGEN) with some modifications (Supplementary Information). HiMEx (three μL of lysate) and ‘conventional’ (1 ng of DNA) extractions were analysed through amplicon sequencing using universal primer set that amplifies prokaryotic 16S, chloroplast 16S, and 18S rRNA genes simultaneously [4]. Almost all HiMEx lysates were successfully amplified, while blanks showed no amplification. We also analysed mock communities with 10× fewer cells than our lowest seawater sample, which performed similarly well (Fig S6). Although intensity of amplification varied, after normalization and pooling, sequence reads were similar between conventional and HiMEx (Tab. S3). Between the methods, similar taxonomic diversity and relative abundances were observed across prokaryotic, microalgae and microbial eukaryotes (Fig. 1A) and community composition of samples at a given timepoint for all DNA extraction methods and volumes were statistically similar based on PERMANOVA results (Tab. S4. A-C), with similar patterns of clustering observed across all domains of microbial communities (Fig. 1B). Replicate variability increased for microbial eukaryotes compared to prokaryotes specifically in microscales which is likely due to comparatively increased stochasticity of eukaryotic cells within given microvolumes samples due to their lower overall cell abundances (Tab. S5). Across all conventional and HiMEx samples the main driver of alpha diversity was time with no significant impact on diversity across extraction methods (Fig. 1C, Tab. S6. A-F), demonstrating HiMEx ability to detect subtle ecological differences over time. Overall, the FTPIG method produced the most consistent and comparable results to conventional extraction (Tab. S4A-C, Tab. S5). The main compositional differences between these two methods were driven mainly by the higher relative abundances of oligotrophic, small, free-living bacteria like *Pelagibacter and* SAR86 in FTPIG versus potentially copiotrophic, particle attached Flavobacteria and *Luminiphilus* [5, 6] compared to the conventional method (Supplementary Information) (Tab. S7). Despite the strong clustering between conventional and HiMEx these differences could reflect either variation in extraction efficiency, or differences between whole seawater and 0.2 μm filtered fraction of seawater. For instance, *Pelagibacter* as an ultramicrobacterium can potentially pass through 0.2-μm-pore-size filters, and has already been reported for possibility of being undersampled in 0.2–1□μm size fraction metagenomes [7–9].

**Fig. 1.**
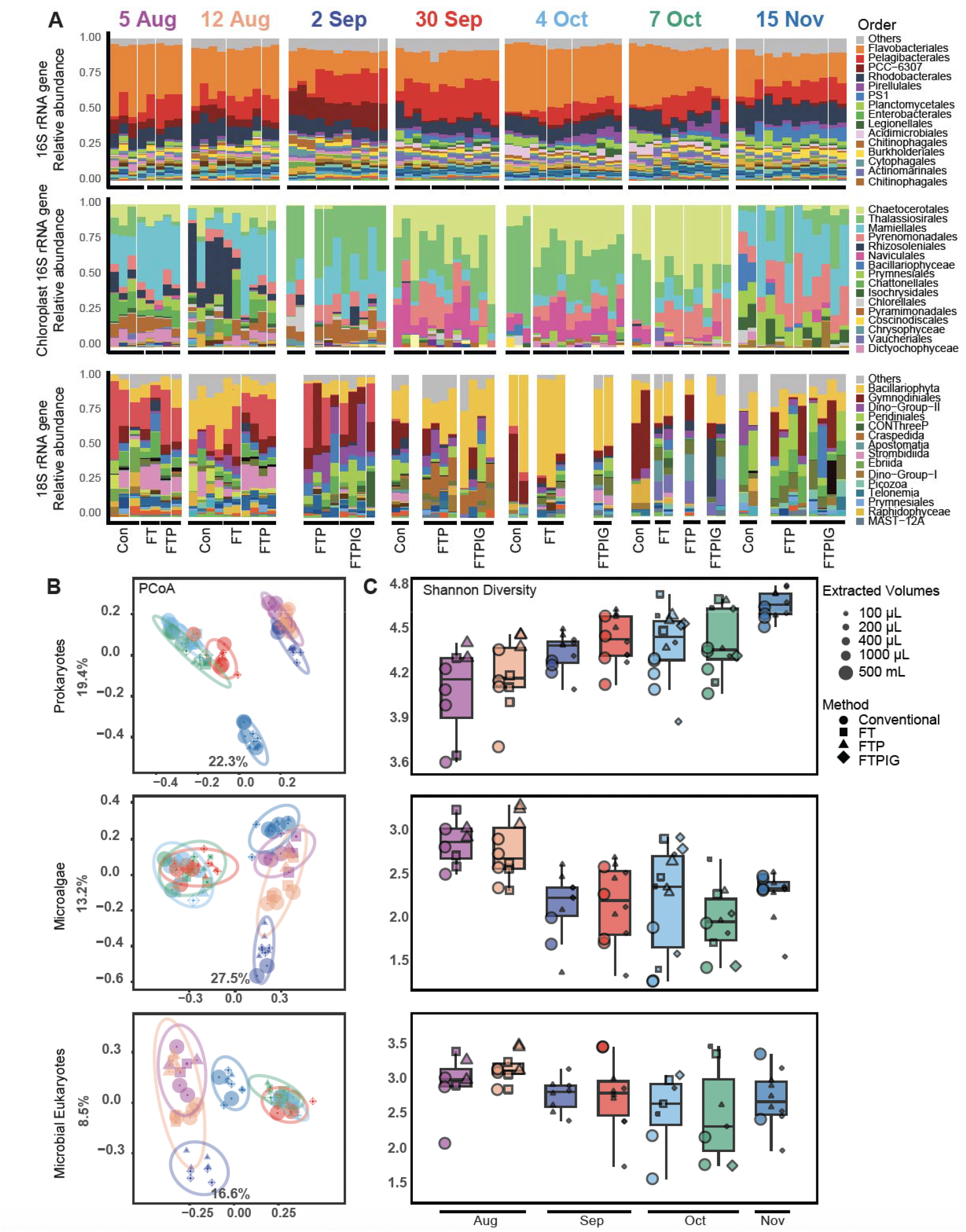
Community compositions via HiMEx and ‘conventional’ 500-mL filter-based DNA extraction are highly concordant. A.) Microbial community composition for prokaryotes, chloroplasts, and microbial eukaryotes consolidated at the Order level, top 30 Orders are coloured in the barplots and the rest combined into “Other”. Top 15 Orders are labelled next to their colour codes (complete colour code presented in Fig. S2). B) PCoA of prokaryotes, chloroplast, and microbial eukaryotes is shown where colours represent different timepoints, shapes indicate the extraction methods, and the size of the points reflects the extraction volumes. The ellipses represent 95% confidence intervals around the clusters. C.) Shannon diversity across conventional and HiMEx showing an increasing trend in prokaryotes overtime from summer to winter vs a decreasing trend in microbial eukaryotes, perhaps detecting a seasonal transition. Linear Mixed effect Models (LMM) were used to assess the effect of different extraction methods and volumes on Shannon diversity while accounting for the fix effect of sampling timepoints (mainly driven by temperature, monthly periods and salinity, Tab. S4. A-C), which showed similar and comparable results between HiMEx and conventional extraction (Tab. S6).

After establishing that recovery of community structure is highly similar across cellular communities between conventional and HiMEx approach, we sought to also establish its utility for shotgun metagenomics through a microcosm phytoplankton-enrichment experiment (Fig. 2A). Using the highest performing HiMEx (FTPIG), we applied a bead-based transposome approach (Hackflex) [10] for both seawater (1 ng of conventional DNA extraction) and manipulated seawater (10 μL of microcosm samples after a 0.8X Ampure bead clean-up) (Supplementary Information). The bead-purification was not strictly required (Fig. S3), but nevertheless served to increase input and fragment size for the metagenomic sequencing (at marginally increased expense). All samples from both conventional extraction and HiMEx yielded similarly high-quality assemblies in terms of overall assembly metrics (Fig 2B), MAG-quality and quantity (Fig. 2C). In terms of MAG diversity, each method recovered similar overall taxa (Fig. S4), though the HiMEx recovered more representative MAGs (i.e., following dereplication of all MAGs), consistent with the overall larger number and variety of sample types (Fig. 2D). Meanwhile, viral genome (contig) quality and quantity was higher from HiMEx relative to the conventional method, owing most likely to removal of free viruses from the conventional filter-based approach (Fig. 2D).

**Fig. 2.**
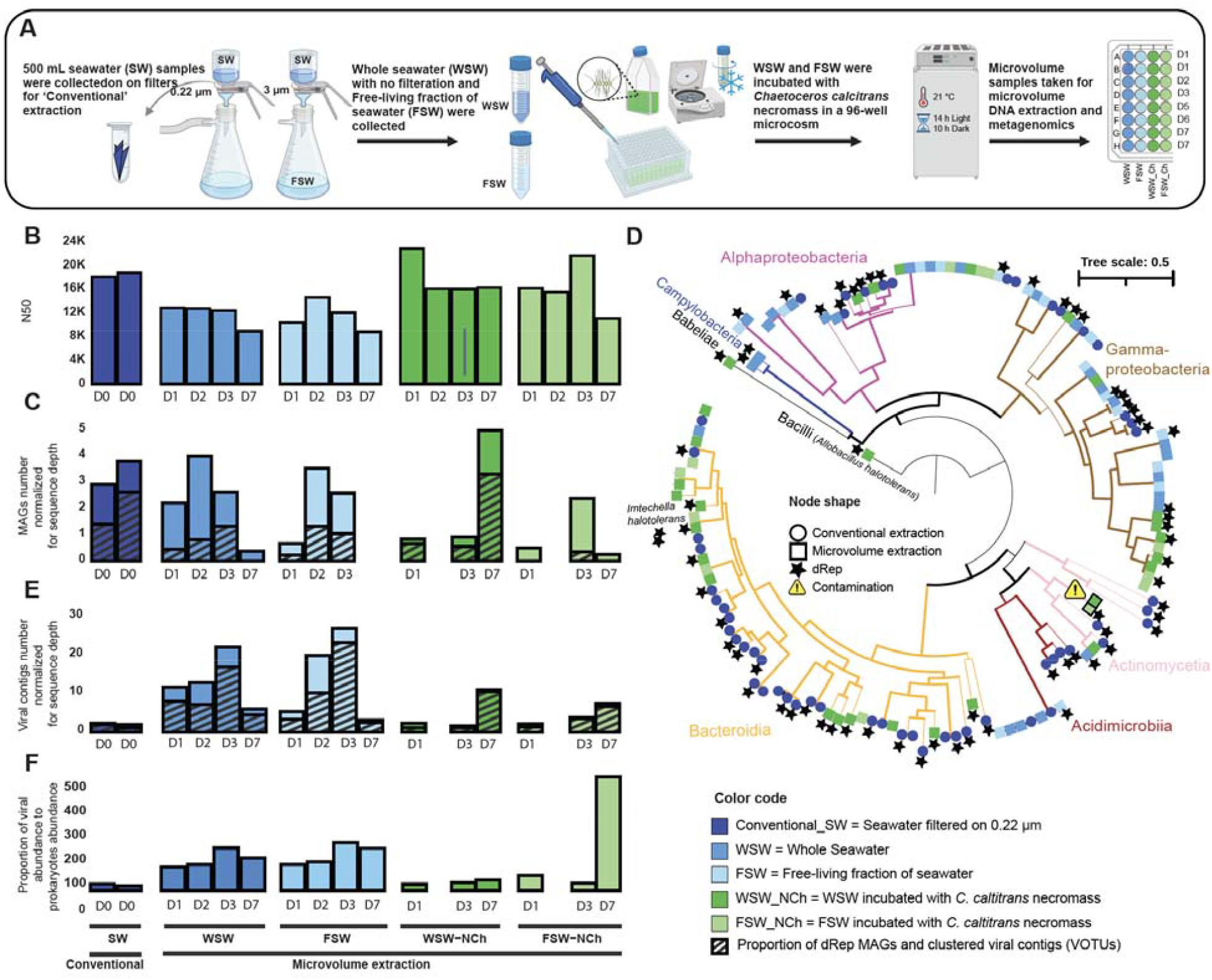
HiMEx extraction yields metagenomic assembly and diversity on par or better than ‘conventional’ filter-based DNA extraction. A) A pilot high-throughput microcosm experiment investigated the impact of microalgal necromass (bloom demise) on multi-domain marine microbial community composition and diversity. Whole seawater and the free-living fraction (<3 μm) of surface Baltic Sea were incubated with *Chaetoceros calcitrans* necromass (obtained after gentle centrifugation and freeze-thaw of cells). Incubation was done in a 96-well microcosm at 21°C, 14 hours light and 10 hours dark for eight days. B) 100-μL daily samples were collected for metagenomic analysis. Overall assembly quality from HiMEx was comparable to that of the conventional extraction based on the total assembly N50. C) Recovery of high- and draft-quality MAGs via conventional and HiMEx approaches were comparable. In order to account for sequencing depth in MAG recovery, their total numbers were divided by total sequencing depth (Tab. S8.A). Across all samples, 138 draft and high quality draft MAGs [19] were produced, with 92 remaining after dereplication (indicated by black hatch patterns, whereby 44 and 48 dereplicated high quality and draft MAGs were recovered from ‘conventional’ and HiMEx samples, respectively (Tab. S9). D) Phylogenomic tree of all MAGs constructed from conventional and HiMEx approach was based on 25 ‘Bacteria_and_Archaea’ single copy gene HMM set protein sequences. Representative MAGs remaining after dereplication whereby MAGs are selected for higher quality, and lower contamination are highlighted with stars. Blanks for HiMEx did not produce any contigs > 5 kb, so MAGs were examined for contaminants list from 16S (Tab. S10) which identified two *Cutibacterium* as potential contaminants, which were excluded from further analyses. In both methods we spiked two bacterial species (*Imtechella halotolerans* and *Allobacillus halotolerans*) equal to an estimated 5% of the cell count to the sample before extraction to help with absolute quantification, however, we only could retrieve high quality MAGs of these spiked bacteria from HiMEx metagenome amplification. E) Excluding proviruses, 465 complete, high- and medium-quality viral contigs were recovered from conventional and HiMEx samples (35 vs 430), with 344 (26 vs 318) VOTUs remaining after clustering (black hatches) (Tab. S11). Similar to bacterial MAGs, to account for sequencing depth, the total number of viral contigs was divided by total sequencing depth (Tab. S8.B). We also obtained 1 vs 38 potentially complete circular viral genomes from conventional extraction and HiMEx, respectively. F) Higher ratios of coverage-based viral to bacterial abundance (VBR) were detected in whole seawater and free-living fraction of seawater compared to the conventional extraction (Tab. S12). Together, these results demonstrate better recovery of viruses from HiMEx vs those recovered from a 0.2 μm filter, in line with expectation that most viruses would go through a 0.2 μm filter.

Our findings indicate that HiMEx produces results for prokaryotes, microalgae, and microbial eukaryotes comparable to a ‘conventional’ filter-based method. Additionally, it offers an innovative, efficient, and cost-effective approach for capturing viral diversity without requiring large sample volumes or the filtration and concentration steps. This method captures small microbes and exocellular DNA that are often missed by filter-based methods. Exocellular DNA is comprised of three main components: protein-encapsidated viruses, exocellular vesicles, and free DNA [11–14], however, their sampling via conventional methods is perhaps thought to require large sample volumes, additional filtration, cost and effort [12, 15]. HiMEx can capture these exocellular DNA as part of whole seawater. However, at present HiMEx cannot separate them from cellular fraction or distinguish between DNA originating from free DNA vs protein-encapsidated viruses, and extracellular vesicles. Hence, a potential consideration is that HiMEx results may include sequences from free-DNA relative to filtration-based methods, along with the increased detection of viruses we observed. Since the proportion of vesicles and free DNA tends to increase with depth [13], this limitation may pose a greater concern in certain environments, such as deeper depth water samples [16]. Free DNA may be addressed by additionally treating the sample with DNase prior to the physical lysis step [17], though this treatment needs further investigation. Additionally, long-term storage of these extractions may be less robust, though has not yet been investigated. Finally, while our analysis demonstrates that HiMEx performs well for surface seawater (with typical prokaryotic abundance of ∼10^6^/mL [18]) and mock communities to 10^4^, (Fig. S5) samples with very low biomass (orders of magnitude lower than surface seawater), amplification and contamination may become an issue.

In conclusion, HiMEx offers an approach for advancing high-throughput, microscale taxonomic and genomic studies in microbial ecology. This approach enables the analysis of hundreds to thousands of samples, supporting large-scale experimentation and microbial diversity assessments in a more cost-effective and sustainable manner.

## Data Availability

Raw reads are available from SRA under bioproject TBD upon publication, or available from the authors upon request. Scripts for analysis and figures are available via https://github.com/Marjan-Ghotbi/HiMEx.

## Author Contributions

MaG and DMN conceptualized the study. Experiments were carried out by MaG. Laboratory work was primarily carried out by MaG, with contributions from MK and DMN. ED carried out filter-based molecular workflow. Field collections were by MaG, MK, ED, and DMN. Data processing and analysis were carried out by MaG, with supervision from DMN and valuable input and guidance regarding statistical analysis from MiG. MaG and DMN wrote the manuscript and all authors contributed to editing the manuscript.

## Acknowledgements

We thank all from Marine Microbial Ecology group at GEOMAR. We thank Jan Muschiol and Kerstin Petersen for general support. Funding was provided by a Young Investigator Grant awarded to DMN.

## Conflict of Interest

The authors declare that they have no conflict of interest.

## Notes

https://github.com/Marjan-Ghotbi/HiMEx

### Competing Interest Statement

The authors have declared no competing interest.

