## Supplementary information for "From Microscale to Microbial Insights: Validating High-Throughput Microvolume Extraction Methods (HiMEx) for Microbial Ecology"

### Materials and Methods

#### Sampling for amplicon analysis

Seawater was collected via bucket from the top meter of Kiel Fjord (54°19.813' N, 10°8.993' E) into a 4 L bottle. Immediately prior to any step of sample processing, the sample bottle was mixed to ensure homogeneity. For HiMEx, a subsample of 50 mL was taken in a centrifuge tube to enable gentle swirling of the water sample before pipetting of each microvolume sample. For conventional method 500 mL was collected on membrane filters (Pall Supor, Fisher Scientific, 0.2 µm, 47 mm; Germany) and for microvolumes, 100 µL-samples were individually pipetted into strips and 1 mL-samples into 2 mL microcentrifuge tubes, and all stored at -80°C until extraction. For high-throughput extraction, 100 µL HiMEx samples were transferred to PCR plates and 1 mL samples were subsampled and transferred in deep-well plates (200, 400, or 1000 µL). However, to facilitate high-throughput analysis, it is recommended to perform sample storage and extraction in the same plate, for further minimizing sample loss and reducing plastic use.

#### Conventional DNA extraction and sequencing for amplicon analysis

Samples consisted of four replicate filters (except for timepoints 92 with three replicates), and three blank replicates including DNA extraction reagents and filters were considered to identify contaminants. For 'conventional', filter samples, *Imtechella halotolerans* and *Allobacillus* *halotolerans* (ZymoBIOMICS Spike-in Control I (High Microbial Load) were added to each sample at roughly 5% (1 to 10%) of the cell count in the sample before extraction (specifically, assuming 1,000,000 cells per mL, 0.25 µL was used for spike-in). A modified extraction was performed based on DNeasy Plant Kit (Qiagen). First, filters were crushed, and then 50 µL of 0.1 and 0.5 mm Carl Roth beads and 750 µL of Lysis buffer AP1 was added, followed by three rounds of freeze and thaw cycles in liquid nitrogen and 65°C water bath, and beat beating in Mixer Mill from Retsch at 30 Hz. Then, lysate was transferred to a 96 well plate, and 45 µL of Proteinase K added followed by incubation for 30 minutes at 55°C. Then, 4uL of RNaseA was added and incubated for an additional 10 minutes at 65°C. 130 µL of Buffer P4 was then added and incubated on ice for 10 minutes, followed by centrifugation to pellet debris. 650 µL of supernatant was added to another 96 well plate and 975 µL of Buffer AW1 added, followed by transfer to a NAB (AcroPrep Advance 96 Filter Plate Nucleic Acid Binding 1 mL well – Pall Corporation). DNA was collected on the column by placing the plate on a vacuum manifold and filtered at less than 250 Hg, until all volume has been collected. Columns were washed 2x with 600 µL of Buffer AW2, and DNA eluted with 3x with 50 µL of pre-heated TE. 1 ng of DNA extract was used as a template for PCR for amplicon library preparations. KAPA HiFi HotStart PCR Kit (Roche, KK2102) was used for amplicon workflow after manufacturer protocol. PCR thermocycling began with an initial denaturation at 95 °C for five minutes, followed by 30 cycles of 20 seconds of denaturation at 98 °C, 15 seconds of annealing at 55 °C, and 1 minute of extension at 72 °C. A final extension step was performed at 72 °C for five minutes. The resulting PCR products were purified and size-selected using 0.8× (vol:vol) Agencourt AMPure XP magnetic beads (Beckman Coulter). Samples were pooled at approximately equimolar concentrations after quantification by Quant-iT PicoGreen dsDNA Assay Kit. The pooled library underwent an additional purification step using 0.8× (vol:vol) AMPure beads before sequencing. Blank samples for extraction were included in amplification and along with PCR blanks were incorporated into the final sequencing libraries.

**HiMEx and sequencing for amplicon analysis**

All tubes, filter tips used during this protocol were sterile and DNA, RNase-Free. Molecular grade water and all reagents used except for the Proteinase-K and DNA polymerase were crosslinked in a UVP crosslinker (CL-3000, Analytik Jena) on the highest energy setting for 1 hr prior to use [1]. The HiMex was performed in a clean bench. Prior to sample processing, the workspace and clean bench were wiped with bleach (10%) and ETOH (70%). Samples were melted on ice and vortexed.

**Physical Lysis**

Samples were heat-shocked for five cycles of freeze-thaw (FT), via rapid alternation between liquid nitrogen (-196 °C) and a 65 °C water bath (Fig. S1). Care was taken not to completely submerge the plate into liquid nitrogen nor the water bath to avoid contamination or sample loss. Samples were briefly vortexed to ensure thorough mixing. Either no centrifugation, or a brief, low speed spin was used to collect the volume at the bottom of the tube, while avoid to pellet the cells and/or debris. Depending on the sample volume a thermoblock or thermocycler was used to heat samples at 99 degree for 10 minutes. Finally, samples were briefly vortexed to mix thoroughly.

**Chemical Lysis**

**FTP treatment (freeze-thaw, plus proteinase K)**

Samples were similarly heat shocked for five cycles of freeze-thaw. Then, 5% v/v of Proteinase K (Qiagen, 19133) was added to each sample except for 1000-μL samples that received 2.5%. Samples were incubated at 55 °C for 20 minutes. Then samples were briefly vortexed to mix and centrifuged very briefly at low speed to collect the reagents. Depending on the sample volume thermoblock or thermocycler was used to heat samples to 99 °C for 10 minutes to deactivate Proteinase K. Finally, samples were briefly vortexed to mix (Fig. S1).

**FTPIG treatment (freeze-thaw, plus proteinase K and IGEPAL)**

Prior to physical lysis step, IGEPAL 20% was prepared fresh by adding PBS (Phosphate-buffered saline) to IGEPAL® CA-630 (Sigma-Aldrich, I3021). Then 1% v/v of IGEPAL 20% was added to each sample (for instance 1 μL of IGEPAL 20% was added to the 100 μL sample). Samples were heat shocked for five cycles of freeze-thaw, and were vortexed short to ensure mixing. 5% v/v of Proteinase K was added to each sample except for 1000-μL samples that received 2.5%. Samples were incubated at 55 °C for 20 minutes. Then they were vortexed short to mix and collect the reagents. Samples were heated to 99 degree for 10 minutes to deactivate Proteinase K. Finally, samples were briefly vortexed to mix (Fig. S1).

For all three HiMEx methods, 3 μL of the lysate was used directly as a template for amplification without any purification steps. PCR reactions were conducted using the KAPA HiFi HotStart PCR Kit (Roche, KK2102) according to manufacturer protocol plus 5% DMSO (Phusion® Hot Start Flex, New England Biolabs). PCR thermocycling began with an initial denaturation at 95 °C for five minutes, followed by 38 cycles of 20 seconds of denaturation at 98 °C, 15 seconds of annealing at 55 °C, and one minute of extension at 72 °C. A final extension step was performed at 72 °C for five minutes. The resulting PCR products were purified and size-selected using 0.8× (vol:vol) AMPure beads. Blank samples for different steps of extraction (mostly in triplicates) were included in amplification and along with PCR blanks were incorporated into the final sequencing libraries. Samples were pooled at approximately equimolar concentrations after quantification by Quant-iT PicoGreen dsDNA Assay Kit. The pooled library underwent an additional purification step using 0.8× (vol:vol) magnetic beads before sequencing. Amplicon libraries were sequenced on Illumina MiSeq (PE300).

**Testing HiMEx on standard microbial community**

Beyond comparing marine microbial communities, we also evaluated potential extraction biases using a cellular mock community primarily composed of human microbiome bacteria with varying cellular abundances. Although all taxa were recovered using HiMEx (FTPIG), there was a noticeable bias against Gram-positive bacteria (Fig. S6) consistent with known biases from widely used extraction kits (https://www.zymoresearch.de/products/zymobiomics-microbial-community-standard). While this is an important consideration, the majority of marine microbial taxa are derived from Gram-negative lineages. Thus, given that Gram-positive bacteria may generally be more challenging to extract than marine bacteria (and yet were still recovered). these results, along with our results from the marine community communities are encouraging. This test also helped to ensure low biomass detection by our HiMEx (FTPIG) extraction. The ZymoBIOMICS Spike-in Control I (High Microbial Load) were added to some sample before extraction in different proportions to estimate detection threshold of the HiMex that were removed from ASV table before visualization.

**Amplicon bioinformatic analysis**

Multiple extraction and PCR controls in addition to a mock community and spiked-in bacteria were included as internal controls and processed identically to environmental samples. This ensured comparability across sequencing runs while minimizing bias and contamination. After inspection, spike-in controls were removed from further analysis. A dual-index sequencing approach was implemented to specifically targets the V4-V5 hypervariable regions of the 16S/18S rRNA genes. The forward primer is I5-8 bp index-XF-YF-515Y, where I5 is Illumina adapter (AATGATACGGCGACCACCGAGATCTACAC), XF is a sequencing primer ACACTCTTTCCCTACACGACGCTCTTCCGATCT, YF- is a 7 'N' 'heterogeneity spacer' meant to increase sequence diversity in the initial Illumina base reads, and the 515Y F primer (GTGYCAGCMGCCGCGGTAA). The reverse primer is I7-8 bp index-XR-YR-926R (with sequences of I7 (CAAGCAGAAGACGGCATACGAGAT), XR is sequencing primer (GTGACTGGAGTTCAGACGTGTGCTCTTCCGATCT), YR is a 7 'N' heterogeneity spacer, and 926R (CCGYCAATTYMTTTRAGTTT). Demultiplexed amplicon sequences were trimmed with cutadapt v4.4 [2] implemented in QIIME2 v2024.10 [3], discarding any sequence pairs not containing the forward or reverse primer (error rate set to 0.2). Amplicon sequences were then split into 16S and 18S pools using bbsplit.sh [4] from the bbtools v4.4 (http://sourceforge.net/projects/bbmap/) against curated 16S/18S databases derived from SILVA 132 [5] and PR2 [6]. The 16S and 18S amplicons were then analysed in parallel to amplicon sequence variants (ASVs) using DADA2 [7] implemented in QIIME2. 16S ASVs were classified with qiime2 classify-vsearch plugin against the SILVA 138.1 database for chloroplast and mitochondria detection. Subsequently, the Greengenes2 database, version 2022.10, was used for classification for compatibility with the GTDB database and metagenomic analysis. ASVs identified as Mitochondria and unassigned reads were removed. Then, the ASV table was subdivided into prokaryotic 16S ASV table and chloroplast 16S ASV table (including all 16S ASV identified as Chloroplast [8]). Chloroplast ASVs were further classified against PhytoRef database87 [9], and finally 19 ASVs identified as order "Embryophyceae", with 474 reads were removed from Chloroplast ASVs to only keep microalgae. 18S ASVs were assigned against the PR2 database. Decontamination analysis was first performed using "prevalence" method of Decontam R package v1.18.0 [10], via the function "isContaminant" which selects contaminant ASVs based on the prevalence (5%) of each sequence feature in true samples compared to the prevalence in negatives and controls (Tab. S10). Hence ASVs representing > 5% of each extraction blank were eliminated from the relevant extraction type. A final decision regarding the inclusion or exclusion of each contaminant ASV was made following a manual inspection and comparison with distributions in GBIF (www.gbif.org/occurrence/) to ensure accuracy based on where these taxa were commonly observed in marine waters and considered non-contaminant. Subsequent to decontamination and removal of blank samples and samples below 1,000 for 16S rRNA and below and 100 reads for chloroplast and 18S rRNA, the total reads per sample varied between 1,000 to 83,675 reads/sample for 16S rRNA, and between 100 to 28,502 for chloroplast and 100 to 21,953 for 18S rRNA. A final number of 79, 76, and 59 remaining samples for 16S rRNA, chloroplast and 18S rRNA were rarefied at 4,000, 100, and 100 sequence depth for prokaryotes, chloroplast and microbial eukaryotes, respectively. These thresholds were set after a thorough check of rarefaction curves to ensure sufficient sequencing depth for reliable community and diversity estimates.

**Amplicon Statistical analysis**

**Assessing microbial community and diversity analysis**

Bray-Curtis distances computed from the rarefied amplicon abundance tables were used to test for homogeneity of dispersion (beta dispersion), through vegan package [11] with 9,999 permutations to determine the variability between replicates depending on extraction types or volumes Generally variability between replicates increased in microvolumes and in FT and FTP Extraction methods (Tab. S5). Then, microbial community variation at different timepoints was assessed using permutational analysis of variance (PERMANOVA) with obtained Bray-Curtis distance matrices. Since some timepoints were testing only three extraction methods, to have balance designs, the dataset was analysed in three subsets with extraction method types equally distributed (Tab. S4.A-C). Surface seawater samples were significantly different at different timepoint. Temperature, salinity and time (month of sampling) and their interactions could explain 0.64, 0.57, and 0.43 of variation in the prokaryotic, chloroplast and microbial eukaryotes datasets, respectively (Tab. S4. A-C). To determine if statistical differences existed at the taxonomic levels between extraction methods and volumes, (PERMANOVA) was computed using the pairwise.adonis function on the subsets of rarefied amplicon abundance tables correcting for the impact of different timepoints as strata, and the *p* value was adjusted throughout (p.adjust = "Holm").

Although PERMANOVA did not show significant difference between conventional and microvolume extractions (Tab. S4. A-C), DESeq2 [12] embedded in DspikeIn R package [13] was used to identify differentially abundant ASVs between conventional vs. FTPIG treatment of HiMEx (with most similar results to conventional extraction) method. In the prokaryotic dataset higher enrichment of *Pelagibacter* was the primary factor driving differences between HiMEx and conventional extraction (Tab. S7). *Pelagibacter*, the most dominant bacteria in the ocean, is classified as an ultramicrobacterium, with a cell volume of less than 0.1 $\mu m^3$ [14], which can potentially pass through 0.2-μm-pore-size filters [15]. In deeper oceanic areas (75 to 500 m) the *Pelagibacter*-like DNA has been reported to dominate the vesicle fractions [16], complicating the interpretation of whether differential abundance reflects cellular DNA or extracellular DNA. However, this concern is likely minimal for surface seawater samples [16].

Linear mixed models (LMM) from the R package lme4 [17] as described in Ghotbi *et. al* [18] were used to estimate the comparative and interactive effects of extraction methods and volumes on prokaryotes, microalgal and microbial eukaryotes diversity (Shannon indices). Time-point (KFTno, Kiel Fjord Time series number, Table S6) was included as a fixed effect to account for the impact of environmental variables which were the main drivers of community clustering). In addition, either extraction method or volume was included also as a fixed effect to test for any additional methodological impact (Table S6). These parameters were also included as random effects to assess their contribution to the partitioning of variance. The ggplot2 [19] and DspikeIn R packages [13] were used for visualization of data.

**Microcosm experimental manipulation and sampling for metagenomic analysis**

A pilot high-throughput microcosm experiment was conducted to examine how bloom demise (phytoplankton necromass) influences the composition and diversity of marine microbial communities across domains and viruses. Surface water from the Baltic Sea, including both whole seawater and its free-living fraction, was kept as seawater control and also incubated with necromass derived from axenic cultures of *Chaetoceros calcitrans*. The *C. calcitrans* cell concentrate was prepared by gentle centrifugation of cells at the end of the exponential phase. After elution in fresh media cellular abundance was $8.29 \times 10^6$ cells/mL (via Gallios flow cytometry). Volumes of 800 µL was added to a deep well plate and subjected to freeze-thaw to obtain *C. calcitrans* necromass. Then 1200 µL of either 3-µm filtered or whole seawater was added to the necromass achieving a final concentration of 3.31 × $10^6$ cells/mL. The deep well plate was incubated at 21 °C under a 14:10 hour light-dark cycle for eight days. Samples of 100 µL were collected daily from different treatments and control samples for metagenomic sequencing. For validation of HiMEx, samples collected from microcosm experiment were compared with initially collected 500 mL water sample which underwent filter-based conventional extraction. ZymoBIOMICS Spike-in Control I were also added to each sample at roughly 5% (1 to 10%) of the estimated cell count in the sample before extraction.

**Sequencing for metagenome libraries**

Sequencing libraries for metagenomes, were prepared using one ng input with 12 to 15 amplification cycles vs. 10 µL lysate with 16 cycles were used for conventional and HiMEx, respectively. Briefly, DNA was tagmented in 50 µL reactions, amplified using the Phusion® Hot Start Flex DNA Polymerase (M0535, New England Biolabs) protocol, excluding DMSO according to the Hackflex protocol [20]. Forward and reverse primers included P5 or P7 adapters, 8-base-indexes, and partial overhangs matching to the transposome adapter TCGTCGGCAGCGTC or GTCTCGTGGGCTCGG, respectively. PCR began with an initial denaturation at 98 °C for 30 seconds, followed by 12 to 15 cycles of 10 seconds at 98 °C for conventional vs 16 cycles for HiMEx, 30 seconds at 62 °C, and 30 seconds at 72 °C, with a final extension at 72 °C for 5 minutes. After amplification, products underwent 0.6× size selection using AMPure XP magnetic beads, followed by bioanalyzer evaluation and an additional 0.6× bead clean-up to remove short DNA fragments. Blank samples were included in all PCR protocols and incorporated into the final sequencing libraries. Metagenome libraries were sequenced on an Illumina NovaSeq 6000 (2 x 150 bp paired reads).

**Metagenome bioinformatic analysis**

Raw sequences were quality-controlled using Trimmomatic (v0.39) [21] to remove adapter sequences, low-quality bases, and short reads. Specifically, reads with a Phred score below 30 were trimmed using a sliding window approach (50:30), and reads shorter than 50 bp were discarded. Post-trimming, read quality was assessed using FastQC [22] to ensure the removal of low-quality regions and adapter contamination. High-quality paired-end and unpaired reads were assembled using SPAdes (v3.11.1) [23], applying the following k-mer values: 21, 33, 55, 77, 99, and 127, with the option –sc. Assemblies were quality checked using MetaQUAST [24] (Fig. 2). Contigs shorter than 5,000 bp were excluded from downstream analyses. Trimmed reads were mapped back to the assembled contigs using Bowtie2 (v2.3.4.3) [25]. To recover metagenome-assembled genomes (MAGs), three binning algorithms were utilized: MaxBin2 [26], MetaBAT2 [27], and CONCOCT [28]. The outputs from the three algorithms were consolidated using DASTool [29] to generate final high-quality bins for each sample. The quality of the resulting bins was assessed using CheckM (v1.1.2) [30] to estimate genome completeness and contamination. Bins were classified as high-quality draft (>90% completeness) or draft (90% completeness) only if contamination levels were below 5%. dRep (v2.6.0) [31] was used for dereplication, removing redundant bins and enabling exploration of how many not redundant and representative genomes were obtained from each extraction method. For decontamination, blank samples were included in and incorporated into the final sequencing libraries. However, after metagenomic analysis workflow blank samples did not yield contigs of at least 5kb (our lowest contig size threshold).

Contigs (from the contaminant MAG classified as *Cutibacterium*) found in the second timepoint of WSW_Ch and FSW_Ch were removed from analysis (Fig. 2), though they were typically undetected in the samples. All MAGs were classified taxonomically using GTDB-Tk (v2.3.2) [32] and the Genome Taxonomy Database (GTDB) release 214.1. The phylogenomic tree of the recovered MAGs was constructed using gtotree (v1.8.1) [33] with the IQtree (v2.1.11) [34], based on high quality draft and draft genomes. Anvi'o was utilized to calculate coverage data based on the Q2Q3 statistic. To enable this, a contigs database was generated within Anvi'o (v8) with anvi-gen-contigs-database, [35] followed by profiling of the samples via anvi-profile and subsequently merged with anvi-merge. Anvi'o Q2Q3 data was calculated via the anvi-summarize command. Q2Q3 data were normalized by the total number of base pairs in the paired-end reads using a per-gigabase scaling approach. This ensured that abundance values reflected microbial community composition across the samples, and these profiles were utilized in downstream statistical analyses to assess microbial community structure and functional capacity. In addition, an end-to-end detection and annotation of plasmids and viruses from contigs equal or larger than 5,000 bp was done through geNomad (v1.8.1) [36] using default settings. Viral sequence quality and completeness were assessed using CheckV [37] (database: checkv-db-v1.5) in end-to-end mode. After excluding proviruses, complete, high and medium quality viral contigs were selected for further analysis which resulted in 465 viral contigs (Tab. S11). To reduce sequence redundancy and dereplicate them, the viral sequences were clustered using cd-hit (4.8.1) [38] with a sequence identity threshold of 95% and an alignment coverage of 85%, with the longest contig being assigned as the representative sequence (VOTU, Tab. S11). 344 high and medium quality VOTUs remained for further analysis.

**Enumeration of phytoplankton, prokaryotes, and viruses in the seawater samples**

For field samples, at each timepoint, 10 mL of seawater was preserved with 0.25% glutaraldehyde incubated for 10–15 minutes at room temperature, and stored at -80 C. Upon thawing, 2 mL of sample was filtered onto 0.02 μM Anodisc filter (Whatman), and stained with 1:100 SYBR Green I following Patel *et al.* [39]. Anodiscs were mounted on to microscopy slides with 0.2% p-phenylenediamine anti-fade mounting medium and visualized on a Zeiss Axio Imager.Z2 Epifluorescence Microscope. Images were acquired in .czi format and analyzed using Fiji (v2.16.0) [40], a distribution of ImageJ bundled with essential plugins for biological imaging. Obtained images were imported via Bio-Format import option which preserved the embedded metadata, including scaling information (pixel size). Pixel size and image scale was verified using the Set Scale tool. For each sample, seven to 10 fields of view were selected at random across the filter depending on image quality. The area of each field of view was calculated using the Analyze > Measure function. Particle counts were performed using Analyze > Analyze Particles function, following manual signal to noise optimization. Particles were categorized into viral-, bacterial-, and phytoplankton-like groups based on relative size ranges, as defined by Patel et al. [39]. Phytoplankton identification was further validated using the chlorophyll fluorescence channel. Number of enumerated particles across six timepoints ranged from 5,669,647 to 50,362,628 for virus-like particles and from 974,372 to 7,558,153 for prokaryote-like particles and from 292 to 17,963 for phytoplankton-like particles (Tab. S1). Based on these data, the lowest bacterial and viral abundances were observed in early September, while the highest abundances occurred in mid-August. The lowest phytoplankton count was recorded in mid-November.

**Supplementary Tables**

Tab. S1. Values represent averages from 7 to 10 counted fields for each sample

Tab. S2. Amplicon analysis Metadata file

Tab. S3. Statistical output of Kruskal-Wallis test on sequencing depth

Tab. S4-A1. PERMANOVA to compare variability in prokaryotic microbial community over different timepoints of KFT (Kiel Fjord time series) after rarefaction at 4000 sequence depth (79 samples)

Tab. S4.A2. PERMANOVA to compare variability in prokaryotic microbial community among different extraction methods after rarefaction at 100 sequence depth (79 samples)

Tab. S4.A3. PERMANOVA to compare variability in prokaryotic microbial community among different extracted volumes (79 samples)

Tab. S4-B1. PERMANOVA to compare variability in microalgae community over different timepoints of KFT (Kiel Fjord time series) (76 samples)

Tab. S4.B2. PERMANOVA to compare variability in microalgae community among different extraction methods (76 samples)

Tab. S4.B3. PERMANOVA to compare variability in microalgae community among different extracted volumes (76 samples)

Tab. S4-C1- PERMANOVA to compare variability in microbial eukaryotes community over different timepoints of KFT (Kiel Fjord time series) after rarefaction at 100 sequence depth (59 samples)

Tab. S4.B2. PERMANOVA to compare variability in microbial eukaryotes community among different extraction methods (59 samples)

Tab. S4.C3. PERMANOVA to compare variability in microbial eukaryotes community among different extracted volumes (59 samples)

Tab. S5. Permutation test for homogeneity of multivariate dispersions in microbial communities via Tukey multiple comparisons of means 95% family-wise confidence level

Tab. S5. Differential abundance of taxa at the Genus level between "conventional" and "FTPIG" extraction methods. "conventional" is used as the reference condition.

Tab. S6-A. Diversity analysis (α-diversity_Shannon indices) of prokaryotic community

Tab. S6-B. Diversity analysis (α-diversity_shannon indices) of microalgae community

Tab. S6-C. Diversity analysis (α-diversity_Shannon indices) of microbial eukaryotes

Tab. S6-D. Diversity analysis (α-diversity_Shannon indices) of prokaryotic community for different extraction volumes

Tab. S6-E. Diversity analysis (α-diversity_shannon indices) of microalgae community for different extraction volumes

Tab. S6-F. Diversity analysis (α-diversity_Shannon indices) of microbial eukaryotes for different extraction volumes

Tab. S7. Differential abundance of taxa at the Genus level between "conventional" and "FTPIG" extraction methods. "conventional" is used as the reference condition. Only those with padj <0.05 are considered significant.

Tab. S8-A. Number of MAGs vs dereplicated MAGs obtained from conventional vs HiMEx method (High-throughput experiment)

Tab. S8-B. Number of complete, high- and medium- quality viral contigs vs dereplicated ones obtained from conventional vs HiMEx method (High-throughput experiment)

Tab. S9. Number and quality of MAGs obtained from conventional vs HiMEx method (High-throughput experiment)

Tab. S10 Contaminants list based on 5% prevelance of 16S dataset in blank samples after manual control

Tab. S11. Number and quality of viral contigs and VOTUs obtained from conventional vs HiMEx method (High-throughput experiment)

Tab. S12. Ratios of coverage-based viral to bacterial abundance (VBR) in conventional vs HiMEx method (High-throughput experiment)

Tab. S13. ASV table (16S rRNA gene) of standard microbial community with different cell number input. Some samples were spiked with specific cell number of internal control which were removed before analysis and making the plot.

**Supplementary Figures**

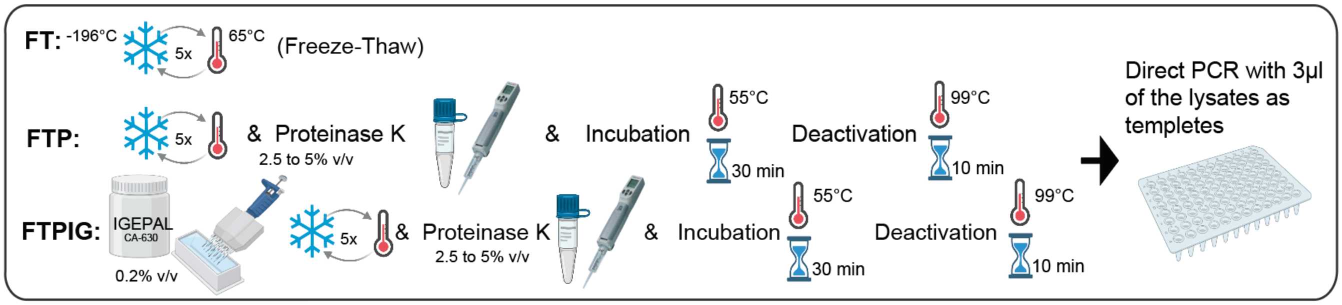


Fig. S1. HiMEx DNA extraction methods for extracting DNA from low volume, low biomass samples. HiMEx relied on either physical extraction through thermal shock (FT) or a combination of physical and chemical extraction which included addition of the lytic enzymes alone to the physical disruption (FTP), or a combination of lytic enzyme and a surfactant (FTPIG). After extraction 3 µL of each lysed sea water was used as a template for amplification via direct PCR.

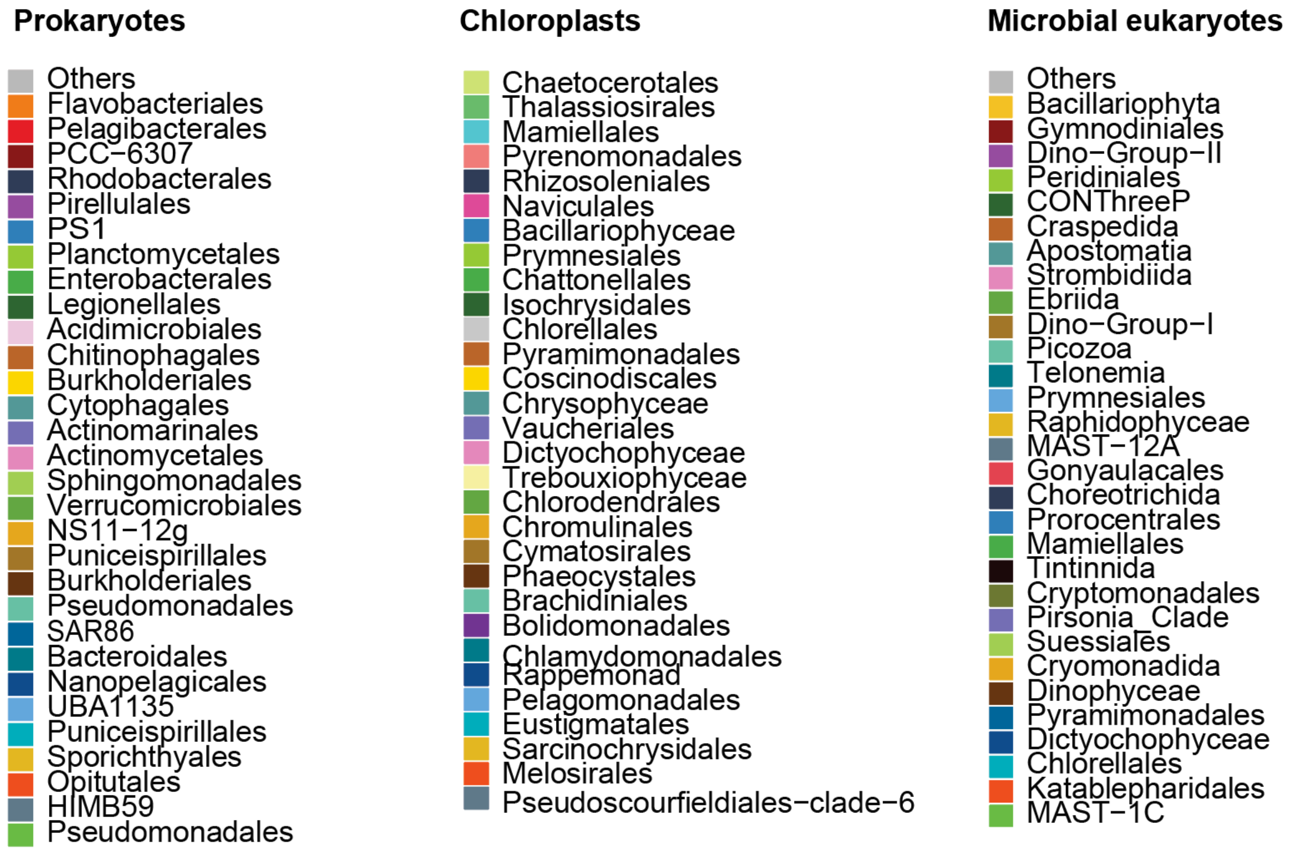


Fig. S2. Color coding for the top 30 orders of microbial community composition in prokaryotes, chloroplasts, and microbial eukaryotes presented in Fig1-A.

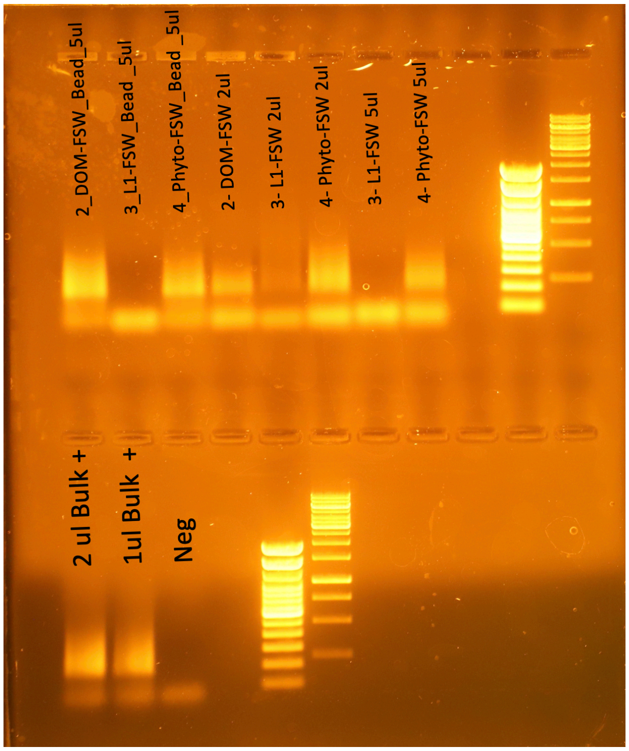


Fig. S3. Gel image showing test samples after HiMEx and amplification using different volumes of lysate (2 to 5 µL) as template for direct PCR and 10 µL using AMPure XP magnetic beads purification (concentrated as 5 µL).

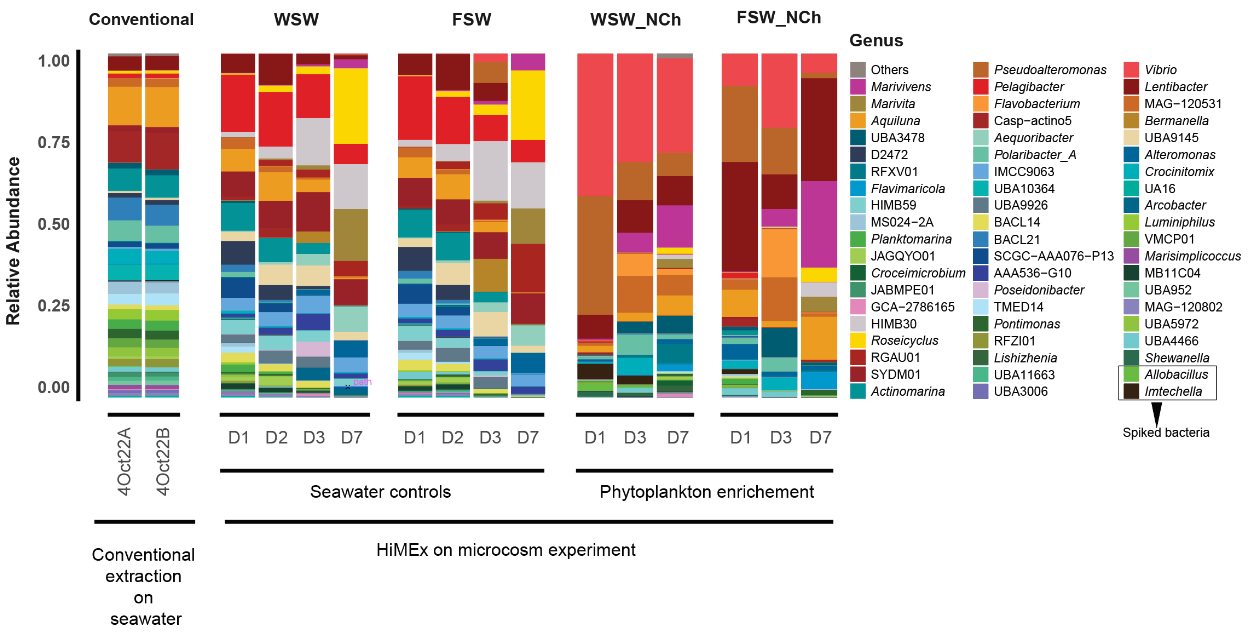


Fig. S4. Overall similar taxonomic diversity recovered by conventional and HiMEx. Incubation of natural bacterial community with phytoplankton necromass clearly increased abundance of *Vibrio*, *Pseudoalteromonas, and Lentibacter* especially in whole seawater samples containing particle-associated bacteria.

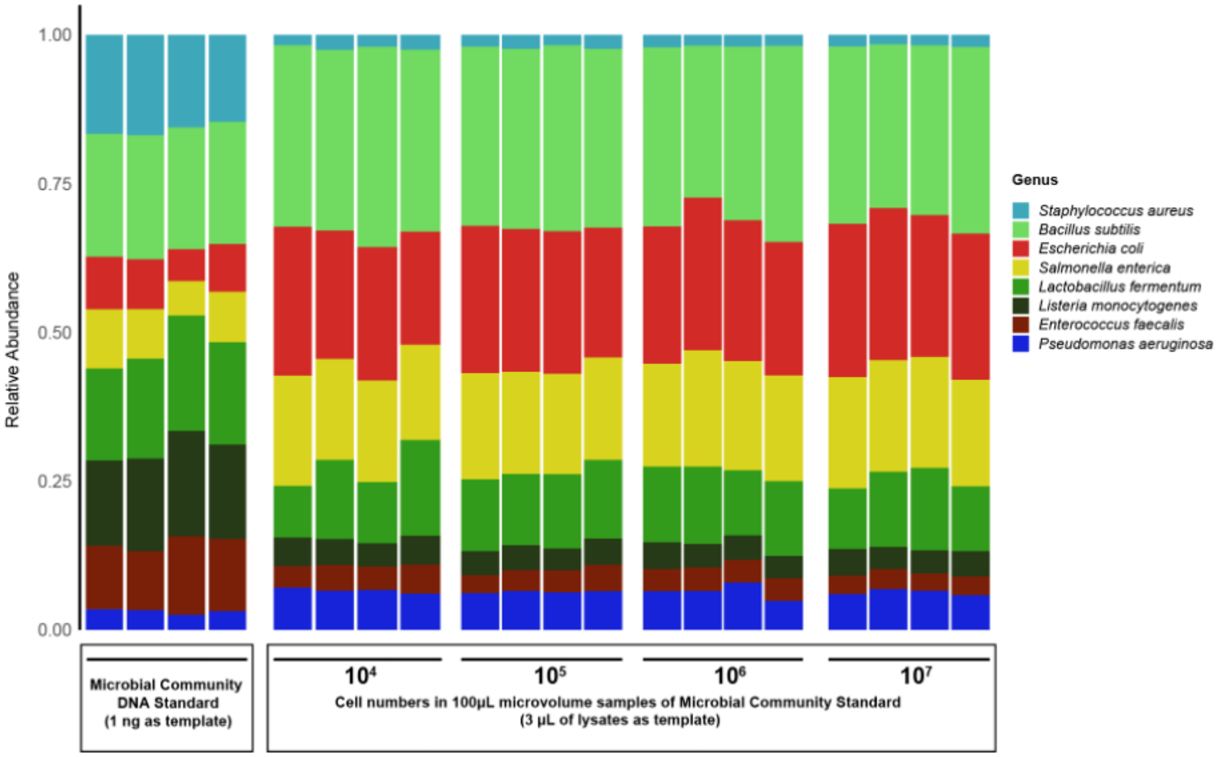


Fig. S5. Barplots showing HiMEx extraction results tested on a gradient of low to medium biomass of standard microbial community (ZymoBIOMICS, D6300), in tandem with DNA standard (ZymoBIOMICS, D6305) (Tab. S13). The cell concentrations selected aimed to assess the sensitivity and ability of HiMEx to amplify cells at lower and higher concentration than typically reported in seawater [41] and our samples (~$10^6$).
